# LPM: a latent probit model to characterize the relationship among complex traits using summary statistics from multiple GWASs and functional annotations

**DOI:** 10.1101/439133

**Authors:** Jingsi Ming, Tao Wang, Can Yang

## Abstract

Much effort has been made toward understanding the genetic architecture of complex traits and diseases. Recent results from genome-wide association studies (GWASs) suggest the importance of regulatory genetic effects and pervasive pleiotropy among complex traits. In this study, we propose a unified statistical approach, aiming to characterize relationship among complex traits, and prioritize risk variants by leveraging regulatory information collected in functional annotations. Specifically, we consider a latent probit model (LPM) to integrate summary-level GWAS data and functional annotations. The developed computational framework not only makes LPM scalable to hundreds of annotations and phenotypes, but also ensures its statistically guaranteed accuracy. Through comprehensive simulation studies, we evaluated LPM’s performance and compared it with related methods. Then we applied it to analyze 44 GWASs with nine genic category annotations and 127 cell-type specific functional annotations. The results demonstrate the benefits of LPM and gain insights of genetic architecture of complex traits. The LPM package is available at https://github.com/mingjingsi/LPM.

## 1 Introduction

In the past decade, genome-wide association studies (GWASs) have been conducted for hundreds of complex phenotypes, including complex diseases and quantitative traits, resulting in the identification of tens of thousands of single-nucleotide polymorphisms (SNPs) associated with one or more complex traits at the genome-wide significance level (1). By exploring these fruitful findings, genetic variants that affect multiple seemly irrelevant traits have been discovered. This phenomenon is known as ‘pleiotropy’ (2). Recently, accumulating studies suggest the pervasiveness of pleiotropy. Pleiotropic effects can be characterized from both local and global perspectives (3). On one hand, localization of pleiotropic risk variants offers more insights on the genetic architecture of human complex traits. For example, a nonsynonymous variant (rs13107325) in the zinc transporter SLC39A8 influences both schizophrenia and Parkinson disease (4); and Ellinghaus *et al.* (5) identified 187 independent multi-disease loci in an analysis of five chronic inflammatory diseases. On the other hand, genetic correlation between two complex traits has been widely explored in recent studies (6), providing a comprehensive view on disease classification (7). Substantial genetic correlations have been revealed among psychiatric disorders, such as the high correlation between schizophrenia and bipolar disorder, moderate correlation between schizophrenia and major depressive disorder, bipolar disorder and major depressive disorder, attention-deficit/hyperactivity disorder and major depressive disorder (8). For autoimmune diseases, primary sclerosing cholangitis and ulcerative colitis, as well as ulcerative colitis and Crohn’s disease, are suggested to have a relatively high genome-wide genetic correlation (9).

The evidence of pervasive pleiotropy not only deepens our understanding of genetic basis underlying complex traits, but also allows the improved statistical power of identification of risk variants by joint analysis of multiple traits. To name a few, joint analysis of schizophrenia and bipolar disorder could significantly improve association mapping power for each of the diseases (10). The power to identify associated variants for systolic blood pressure was increased by considering GWASs of other phenotypes, such as low-density lipoprotein, body mass index and type 1 diabetes mellitus (11).

An increasing number of reports suggest that SNPs with important functional implications can explain more heritability of complex traits (12, 13, 14) and the pattern of enrichment in a specific genic annotation category is consistent across diverse phenotypes (12). For example, SNPs in 5’UTR, exons and 3’UTR of genes are significantly enriched, SNPs in introns are moderately enriched and intergenic SNPs are not enriched in height, schizophrenia and tobacco smoking (12). It is coincidence with the finding that pleiotropic SNPs are more often exonic and less often intergenic compared with non-pleiotropic SNPs (15). Additionally, some cell-type specific functional annotations are also shown to be relevant to complex traits. For example, functional annotations in liver are relevant to lipid-related traits, such as low-density lipoprotein, high-density lipoprotein and total cholesterol (16, 17); functional annotations in immune system are relevant to many autoimmune diseases, such as Crohn’s disease, ulcerative colitis and rheumatoid arthritis (17). Large amounts of functional annotation data have been provided by the Encyclopedia of DNA Elements (ENCODE) project (18) and the NIH Roadmap Epigenomics Mapping Consortium (16).

With the availability of functional annotation data and summary statistics from GWASs on a wide spectrum of phenotypes, we aim to propose a unified framework which can (i) characterize relationship among complex traits, including identifying pleiotropic associations and estimating correlations among traits, (ii) increase the association mapping power for one or more traits, and (iii) investigate the effect of functional annotations. Existing statistical methods based on summary statistics are not able to achieve these aims simultaneously. Methods such as cross-trait LD Score regression (19) and GNOVA (20), provide genetic correlation estimation for pair of traits, but are not able to prioritize GWAS results. In contrast, RiVIERA (21) can prioritize disease-associated variants by joint analysis of summary statistics across multiple traits and epigenomic annotations, but does not measure pleiotropy. Other methods such as GPA (10) and graph-GPA (22), can both infer the relationship among traits and identify causal variants. However, statistical and computational challenges arise as the number of traits increases. GPA assumes a four-group model for the case of two GWASs. The number of groups increases exponentially with the number of traits. Graph-GPA is not able to integrate functional annotations, and its implementation is based on an MCMC algorithm which is time-consuming. Additionally, the relationship among traits inferred by graph-GPA can be hard to interpret in real data analysis, because graph-GPA suggests a graphical model based on an Markov random field which represents a conditional independent structure for genetic relationship among traits and the structure may change when adding or removing some traits.

Here we propose a latent probit model (LPM) to characterize relationship among complex traits by integrating summary statistics from multiple GWASs and functional annotations. To make LPM scalable to millions of SNPs and hundreds of traits, instead of working with a brute-force algorithm to handle all the data simultaneously, we develope an efficient parameter-expanded EM (PX-EM) algorithm for pair-wise analysis and implement a dynamic threading strategy to enhance its parallel property. This pair-wise strategy is guaranteed to give consistent results by our theoretical analysis from the perspective of the composite likelihood approach (23). We conducted comprehensive simulations to evaluate the performance of LPM. Then we analyzed 44 GWASs of complex traits with nine genic category annotations and 127 cell-type specific functional annotations using LPM. The results demonstrate that our method can not only fulfill the three goals (characterizing relationship, prioritizing SNPs and integrating functional annotations) under a unified framework, but also achieve a better performance than conventional methods.

## 2 Materials and methods

### 2.1 GWAS summary statistics and functional annotations

The source of the summary statistics of 44 GWASs is given in Supplementary Table S1. The traits are across a wide range of domains, including psychiatric disorders (e.g. bipolar disorder and schizophrenia), hematopoietic traits (e.g. mean cell haemoglobin concentration and red blood cell count), autoimmune diseases (e.g. Crohn’s disease and ulcerative colitis), lipid-related traits (e.g. HDL and triglycerides) and anthropometric traits (e.g. height and BMI). The genic category annotations were provided by ANNOVAR (24). In this paper, we used nine genic category annoations which include upstream, downstream, exonic, intergenic, intronic, ncRNA_exonic, ncRNA_intronic, UTR’3 and UTR’5, where ncRNA referred to RNA without coding annotation. The cell-type specific functional annotations were generated using epigenetic markers (H3k4me1, H3k4me3, H3k36me3, H3k27me3, H3k9me3, H3k27ac, H3k9ac and DNase I Hypersensitivity) in 127 tissue and cell types from the Epigenomics Roadmap Project. We collected 127 cell-type specific functional annotations from GenoSkylinePlus (25) (http://genocanyon.med.yale.edu/GenoSkyline). We excluded the SNPs in the MHC region (Chromosome 6, 25-35 Mb) to avoid unusually large GWAS signals.

### 2.2 Methods for comparison

#### 2.2.1 GPA

GPA (10) is a statistical approach to prioritizing GWAS results incorporating pleiotropy and annotation. The GPA package can be downloaded from GitHub (https://github.com/dongjunchung/GPA). We followed the package vignette to fit GPA models for pairs of GWASs with and without functional annotations, and implement association mapping and hypothesis testing of pleiotropy. We compared GPA with LPM in both simulations and real data analysis.

#### 2.2.2 graph-GPA

Graph-GPA (22) is a graphical model for prioritizing GWAS results and investigating pleiotropic architecture. The GGPA package can be downloaded from GitHub (https://github.com/dongjunchung/GGPA). We followed the package vignette to fit graph-GPA models for *P*-values of multiple GWASs, implement association mapping and investigate pleiotropic architecture using the default settings. We compared graph-GPA with LPM in both simulations and real data analysis.

#### 2.2.3 RiVIERA

RiVIERA (21) is a Bayesian model for inference of driver variants using summary statistics across multiple traits and epigenomic annotations. It takes linkage disequilibrium (LD) into account and assumes that there is one causal variant per locus. RiVIERA is able to provide 95% credible sets for each locus. The RiVIERA package can be downloaded from GitHub (https://github.com/yueli-compbio/RiVIERA-beta). We followed the package vignette to fit RiVIERA using the default settings. We compared RiVIERA with LPM in simulations.

#### 2.2.4 cross-trait LD Score regression

Cross-trait LD Score regression (19) is a technique for estimating genetic correlations among traits. It can be implemented in LDSC which can be downloaded from GitHub (https://github.com/bulik/ldsc). We followed the ldsc wiki to estimate the genetic correlation for each pair of traits. We filtered GWAS data to HapMap3 SNPs (w_hm3.snplist.bz2) and used the LD Scores (eur_w_ld_chr.tar.bz2) as the independent variable and weights in the LD Score regression. The files can be downloaded from https://data.broadinstitute.org/alkesgroup/LDSCORE. We compared cross-trait LD Score regression with LPM in real data analysis.

### 2.3 Latent probit model (LPM)

Suppose we have the summary statistics (*P*-values) for *M* SNPs in *K* GWASs. In this paper, we use *j* = 1, …, *M* to index SNPs and *k* = 1, …, *K* to index GWAS data sets. For each GWAS, we consider the *P*-values following a two-group model (26), i.e., a mixture of null and non-null distributions, and introduce a latent variable *η_jk_* to indicate which group the *j*-th SNP belongs to for the *k*-th GWAS. Here *η_jk_* = 0 and *η_jk_* = 1 indicate the *j*-th SNP is un-associated (in the null group) and associated (in the non-null group) with the *k*-th trait, respectively. We assume the *P*-values in the *k*-th GWAS to be distributed as

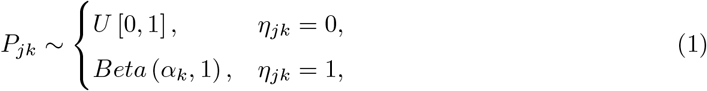

where *U* [0, 1] is the uniform distribution on [0, 1] and *Beta* (*α_k_*, 1) is the beta distribution with the constraint 0 < *α_k_* < 1. This model is designed to capture the pattern that *P*-values from the non-null group have higher density near zero (10, 17).

To adjust the effect of functional annotations and model the relationship among traits, we consider the following latent probit model (LPM):

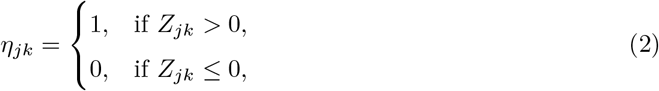

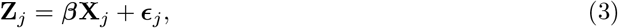

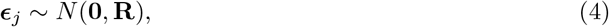

where **Z** ∈ ℝ^*M*×*K*^ is the matrix of latent variables, **X** ∈ ℝ^*M* × (*D*+1)^ is the design matrix of functional annotations, comprised of an intercept and *D* annotations, and *β* ∈ ℝ^*K* × (*D*+1)^ is the matrix of coefficients. For the *j*-th SNP, 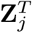, 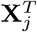 and 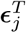 correspond to the *j*-th row of **Z**, **X** and **ϵ**, respectively.

Furthermore, **R** ∈ ℝ^*K*×*K*^ is the correlation matrix measuring the relationship among *K* traits. If the correlation between any two traits exists, the corresponding entry in **R** is expected to differ from 0. We let ***θ*** = {***α***, ***β***, **R**} be the collection of model parameters.

Instead of working with *K* traits simultaneously, we analyze the GWASs in a pair-wise manner based on the composite likelihood approach (see more details in the discussion section). In this case, *K* = 2 and we denote this model as bivariate LPM (bLPM):

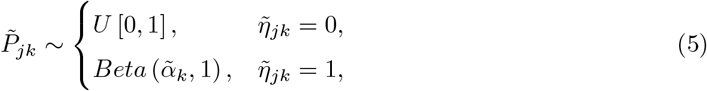

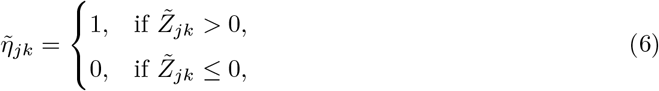

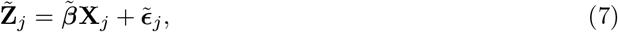

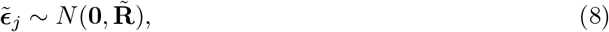

where *k* = 1, 2, *β̃* ∈ ℝ^2×(*D*+1)^ and

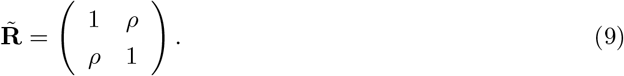

Let ***θ̃*** = {***α̃***, ***β̃***, **R̃**} be the collection of parameters in bLPM. The logarithm of the marginal likelihood can be written as

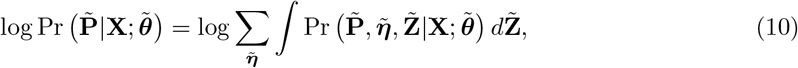

where

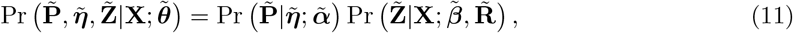

and

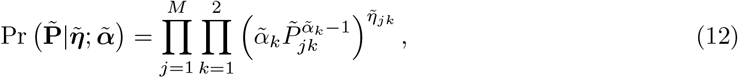

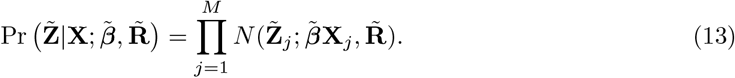

In (12) and (13), we assume conditional independence among *M* SNPs given annotation matrix **X**. Our goal is to find ***θ̃***, which maximizes the marginal likelihood (10) for each pair of GWASs and then obtain an estimate of ***θ*** in LPM. Denote the estimate of ***θ*** as ***θ̂***. We can compute the posterior

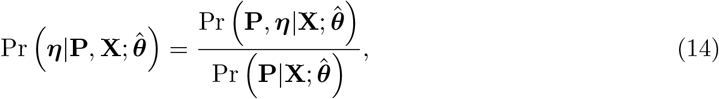

and make statistical inferences on the association of SNPs and relationship among traits.

### 2.4 Algorithm for LPM

The standard EM algorithm converges slowly on LPM. Instead, we propose a PX-EM algorithm, which converges much faster than the standard EM algorithm (27), for parameter estimation and posterior calculation in bLPM. We expand the parameter in bLPM to **Θ** = {***α̃***, ***γ***, **Σ**}. Accordingly, models (7) and (8) are expanded to

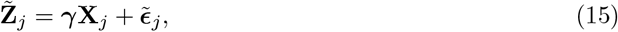

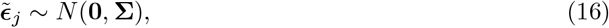

where ***γ*** = **D*β̃***

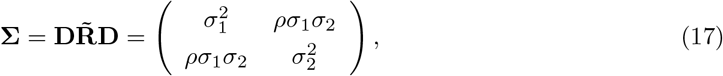

and

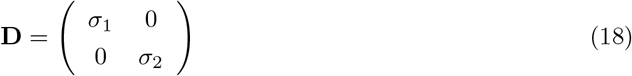

is the auxiliary parameter whose value is fixed at the identity matrix in the original model.

For the expanded model, the complete-data log-likelihood can be written as

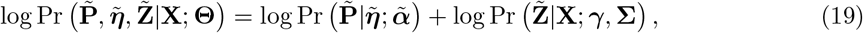

where

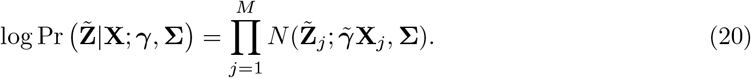

In the PX-E step, the Q function is evaluated as

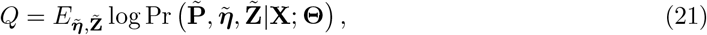

where the expectation is calculated based on the current **Θ** in the original model.

In the PX-M step, we maximize the Q function with respect to **Θ** and obtain the updating equations for each iteration

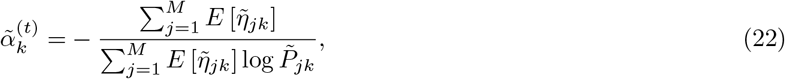

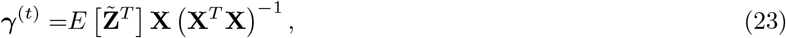

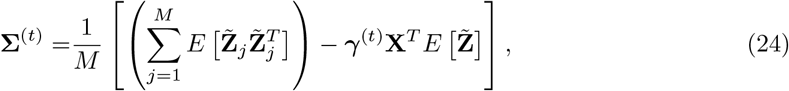

where the superscript (*t*) denotes the *t*-th iteration.

In the reduction step, we obtain the estimates for the original parameters:

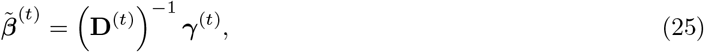

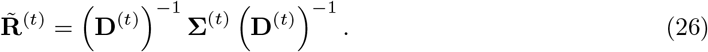

The details of the derivation can be found in the Supplementary Section S1.

If the correlation coefficient *ρ* is zero, we can analyze the traits independently, which provides warm starts for generating our three-stage algorithm for bLPM. In the first stage, we set all the coefficients in ***β̃*** (except the intercept term) and the correlation coefficient *ρ* to be zero, and run an EM algorithm to obtain the estimates for ***α̃*** and ***β̃***_0_. Then in the second stage, we use the estimated parameters as the starting point to obtain ***α̃*** and ***β̃*** using a PX-EM algorithm. Finally, in the third stage, we run the above PX-EM algorithm, using initial parameters those obtained in the second stage, and update ***α̃***, ***β̃*** and *ρ* simultaneously until convergence. Since our algorithm is based on the framework of EM and PX-EM, the likelihood is guaranteed to increase at each iteration. The details of the algorithm are provided in the Supplementary Section S2.

For *K* GWASs, we analyze them pairwisely using the above algorithm and obtain the corresponding estimates ***θ̃̂***. We can implement this procedure parallelly. To obtain the estimates *α̂_k_* and *β̂_k_* for LPM, we use the average over the pairs containing the *k*-th GWAS:

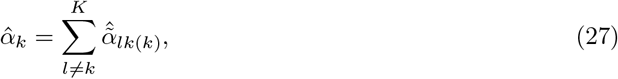

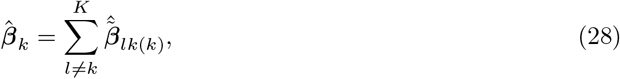

where ***α̃̂**_lk_* and ***β̃̂**_lk_* are the estimate of ***α̃*** and ***β̃*** respectively in bLPM for the *l*-th trait and the *k*-th trait, and (*k*) means the entry for the *k*-th trait. We can also form a matrix **R̂**_*pair*_ using the corresponding estimation *ρ̂*in pairwise analysis. In real data analysis, the number of SNPs *M* is often different in each GWAS due to different genotyping platform and quality control. To avoid losing much information, we allow different *M* in each bLPM. However, since the pairwise analysis is not based on the same data, **R̂**_*pair*_ may not be positive semidefinite, which is required for a correlation matrix. As such, we solve the following optimization problem to obtain the nearest correlation matrix **R̂**:

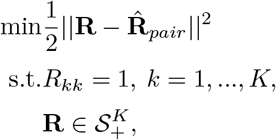

where 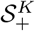 is the cone of positive semidefinite matrices in the space of *K* × *K* symmetric matrices, and || || is the Frobenius norm. This optimization problem can be efficiently solved by a Newton-type method (28).

### 2.5 Inferences based on LPM

#### 2.5.1 Identification of risk SNPs

After we obtain the estimates of parameters in LPM, we are able to prioritize risk SNPs based on the posterior of ***η***, which indicates the strength of association of the SNPs with the traits. If we consider the traits separately, the association mapping of the *j*-th SNP on the *k*-th trait can be inferred from Pr (*η_jk_* = 1|*P_jk_*, **X**). In this case, the relationship among traits is ignored and only the current GWAS data is used.

If two traits are considered, risk SNPs for both the *k*-th trait and the *k′*-th trait can be inferred from

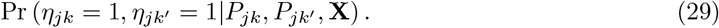

In addition, we can infer the risk SNPs for the *k*-th trait by calculating the marginal posterior

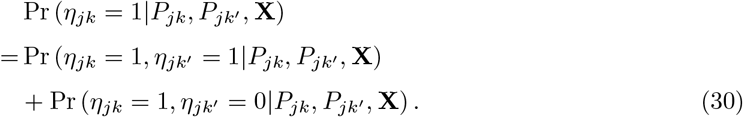

Similarly, we can consider more than two traits, e.g. three traits, and obtain the posterior that the *j*-th SNP is associated with the *k*-th, *k′*-th and *k″*-th trait

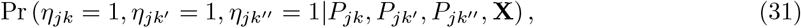

the marginal posterior that the *j*-th SNP is associated with the *k*-th and *k′*-th trait

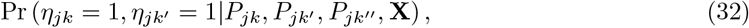

and the marginal posterior that the *j*-th SNP is associated with the *k*-th trait

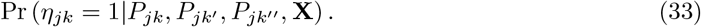

Moreover, we can calculate the local false discovery rate and use the direct posterior probability approach (29) to control the global false discovery rate (FDR). The details are shown in Supplementary Section S3.

#### 2.5.2 Relationship test among traits

We test the relationship between two traits in the pairwise analysis by the hypothesis:

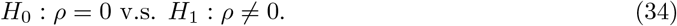

We use the likelihood ratio test. The test statistic is

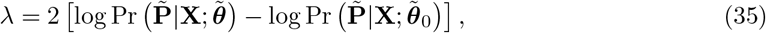

where ***θ̃***_0_ is the parameter estimates under *H*_0_, i.e., the estimates we obtain in the second stage of the algorithm. The probability distribution of *λ* is asymptotically a *χ*^2^ distribution with *df* = 1 under the null.

#### 2.5.3 Hypothesis testing of annotation enrichment

When we integrate functional annotation data, we are interested in the enrichment of annotation for a specific trait. We consider the following test:

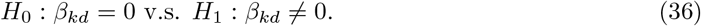

To estimate the standard error of *β̂_kd_* we consider the single trait case. The log-likelihood for the *k*-th trait is

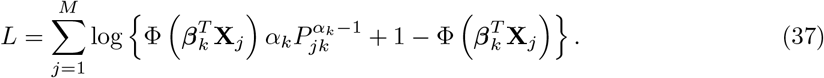

The information matrix can be computed by

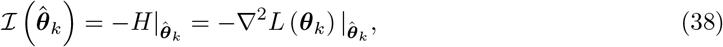

where 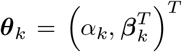 and ***θ̂**_k_* is the estimate in LPM. The details for calculation are shown in Supplementary Section S4.

Then the inverse of observed information matrix provides an estimator of the asymptotic co-variance matrix. The Wald test statistic is

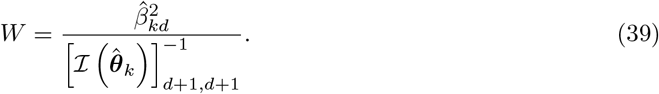

The probability distribution of *W* is approximately a *χ*^2^ distribution with *df* = 1 under the null.

### 2.6 Simulation of eight traits

To evaluate the performance of the proposed LPM, we generated the simulation data for multiple traits using the generative model. The procedure was as follows. We considered eight traits which were divided into three groups: (i) P1, P2 and P3; (ii) P4, P5, P6; and (iii) P7, P8. Correlation existed only within the groups. Specifically, we set the correlation matrix **R** with the corresponding entries *ρ*_12_ = 0.7, *ρ*_13_ = 0.4, *ρ*_23_ = 0.2, *ρ*_45_ = 0.6, *ρ*_46_ = 0.3, *ρ*_56_ = 0.1 and *ρ*_78_ = 0.5 (all the other entries were set to zeros). The relationship among the traits is depicted in Figure 1a. The numbers of SNPs and functional annotations were set to be *M* = 100, 000 and *D* = 5, respectively. First, we generated the design matrix **X** and coefficients *β* of functional annotations. The entries in **X** excluded the intercept were generated from *Bernoulli* (0.2). The entries in first column of *β* were set to be −1 and the other entries were first generated from *N* (0, 1) and then transformed to control the relative signal strengh between annotated part and un-annotated part 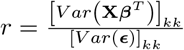 for *k* = 1, …, 8 to be 1. Both **X** and *β* were kept fixed in multiple replications. Then we simulated *η_jk_* according to the multivariate probit model (2)-(4). Finally, we generated *P_jk_* from *U* [0, 1] if *η_jk_* = 0 and *Beta* (*α_k_*, 1) if *η_jk_* = 1 with *α*_1_ = 0.2, *α*_2_ = 0.35, *α*_3_ = 0.5, *α*_4_ = 0.3, *α*_5_ = 0.45, *α*_6_ = 0.55, *α*_7_ = 0.25 and *α*_8_ = 0.4.

**Figure 1:**
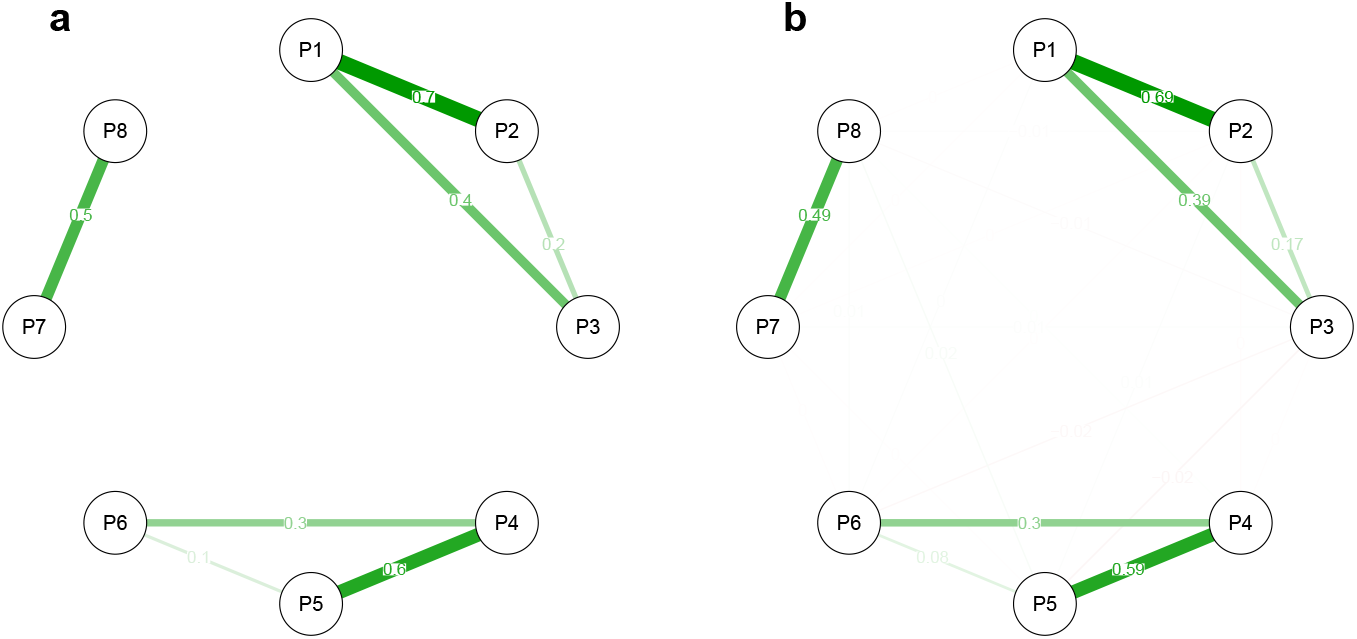
Correlation graphs. **(a)** true graph. **(b)** average estimated correlation graph using LPM. The numbers on the edges and the widths of the edges indicate the correlation between the connected traits. The results are summarized from 50 replications.

## 3 Results

We conducted comprehensive simulation studies to evaluate the performance of the proposed LPM and then applied it to analyze 44 GWASs with nine genic category annotations and 127 cell-type specific functional annotations. In spirit of reproducibility, all simulation codes and real data sets in this study have been made publicly available at https://github.com/mingjingsi/LPM.

### 3.1 Simulations: performance in characterizing the correlations among the traits

We first evaluated the performance of LPM in characterizing the correlations among the traits in our simulations. We simulated summary statistics of eight GWASs and the design matrix of five functional annotations (Materials and methods). Figure 1b shows that the correlation graph is accurately estimated using LPM. We also evaluated the type I error rate and power of LPM for the relationship test among the traits. As shown in Figure 2a, the type I error rates are almost 0 for all the pairs with no correlation and the powers are almost 1 except for two pairs (P2 and P3, P5 and P6) in which cases the correlations are relatively small and the signal strength is relatively weak, i.e., the corresponding *ρ*s are relatively small and *α*s are relatively large. The comparison results of GPA and graph-GPA are shown in Figures 2b and 2c. As the relationship both GPA and graph-GPA measured does not adjust the effect of functional annotations, more significant relationships are detected. If all the functional annotations have no role in the simulation, i.e., **X** only had the intercept term and *β* only had one column in which the entries were set to be −1, the relationship test graphs of LPM, GPA and graph-GPA are similar (see Supplementary Fig. S1).

**Figure 2:**
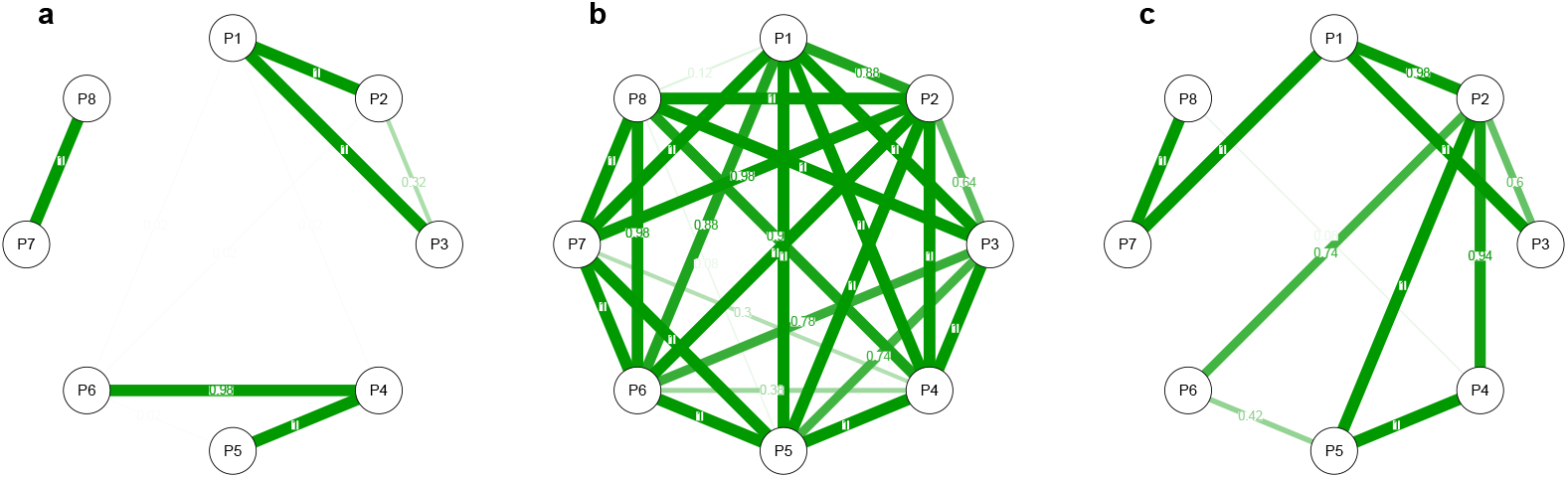
Relationship test graphs of **(a)** LPM, **(b)** GPA, and **(c)** graph-GPA. The numbers on the edges and the widths of the edges indicate the type I error or power of the relationship test for the connected traits. For LPM and GPA, we controlled family-wise error rate at 0.05. The results are summarized from 50 replications.

To provide a better illustration for the performance of LPM, we conducted more simulations which considered only two traits. In this simulation, we set the signal strength of the traits to be the same, i.e., *α*_1_ = *α*_2_ = *α*. We varied *α* in {0.2, 0.4, 0.6} and *r* in {0.25, 1, 4} to obtain the type I error rate, and varied *ρ* in {0, 0.05, 0.1, 0.15, 0.2, 0.25} to obtain the power of LPM for the relationship test between the traits. As shown in Figure 3, the type I error rates are well controlled in all cases and the power increases as *α* decreases and as *ρ* increases. However, we noted that a large relative signal strength *r* could lead to a small power. This is because the correlation resulting from annotations increases as *r* increases and the correlation we aim to estimate becomes relatively smaller.

**Figure 3:**
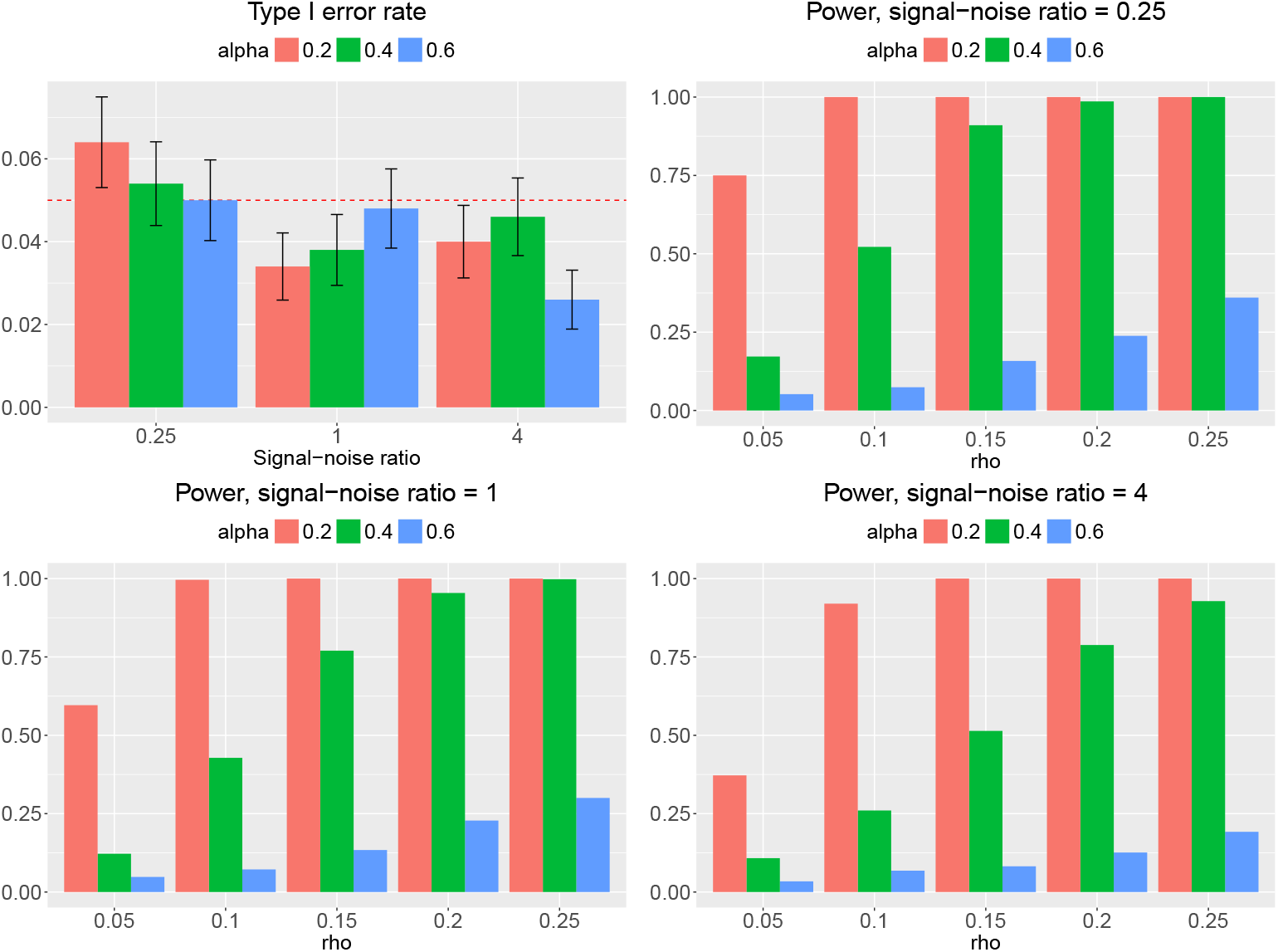
Type I error rate and power of LPM for the relationship test between two traits. The bars represent one standard error. We evaluate type I error rate and power at 0.05 significance level. The results are summarized from 500 replications.

### 3.2 Simulations: performance in the identification of risk SNPs for one or multiple traits

We further evaluated the performance of LPM in the identification of risk SNPs for one or multiple traits in the simulation of eight traits (Materials and methods).

To identify risk SNPs for one specific trait, we consider three cases (i) separate analysis of the target trait, (ii) joint analysis of the target trait with another trait, and (iii) joint analysis of the target trait with other two traits, using LPM. If the integrated traits are correlated with the target trait, the power to identify risk SNPs is expected to increase in joint analysis. We compared LPM with GPA under these three cases in terms of their empirical FDR and AUC. The results for P1 are shown in Figure 4 (results for other traits are given in Supplementary Figs S2-S8). The empirical FDRs of LPM are indeed controlled at the nominal level. However, the FDRs of GPA are inflated in some cases when the GWAS signal is relatively weak and are conservative when the GWAS signal is relatively strong. Moreover, LPM outperformed GPA for all the cases in terms of AUC. As expected, the AUC of LPM increases when correlated traits are integrated. For example, as shown in Figure 4b, the power to identify risk SNPs for P1 increases when the correlated traits (P2 and P3) are jointly analyzed. Specifically, integrating traits with high correlation with the target trait could result in a better improvement of AUC (simulations are described in Supplementary Section S6.3). We also compared LPM with RiVIERA. The details of simulation are given in Supplementary Section S6.4. As shown in Supplementary Fig. S10, LPM achieved a better performance compared to RiVIERA.

**Figure 4:**
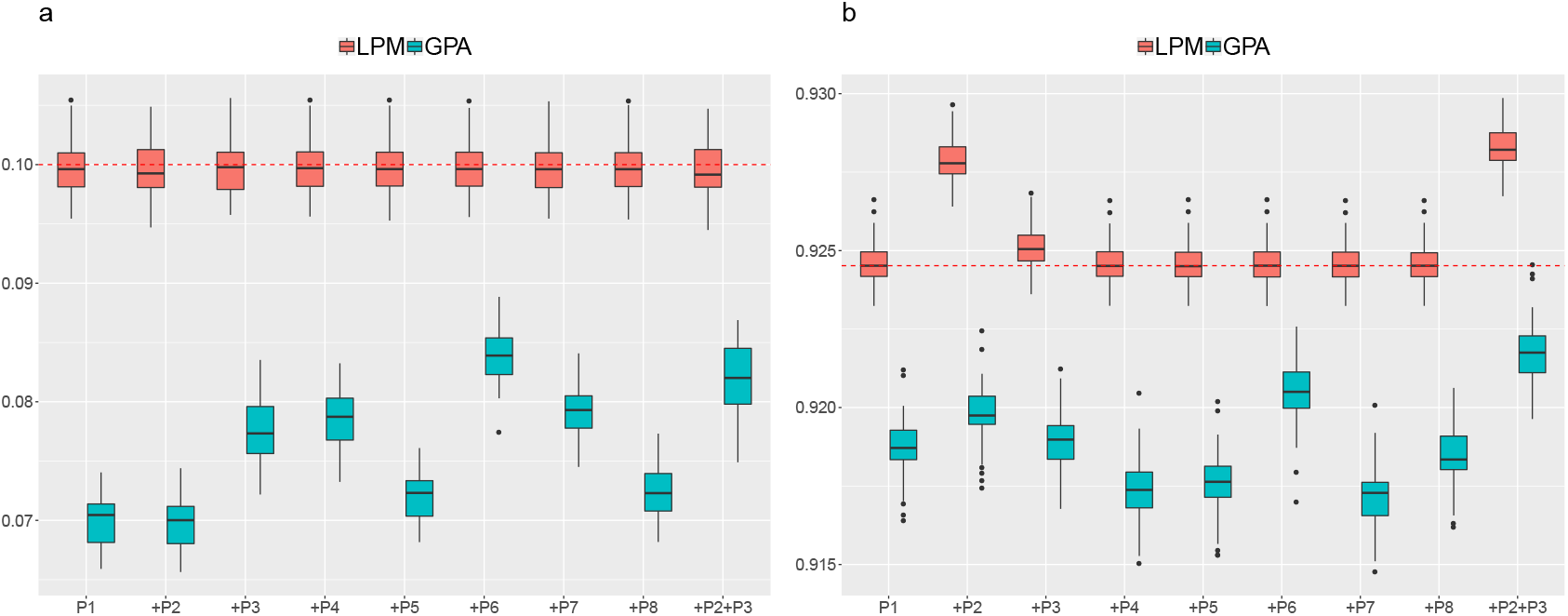
**(a)** FDR and **(b)** AUC of LPM and GPA for identification of risk SNPs for P1. The text in the x-axis indicates which traits are used in analysis, e.g., P1 indicates separate analysis using only trait P1, +P2 indicates joint analysis of P1 and P2. We controlled global FDR at 0.1 to evaluate empirical FDR. The red horizontal line in **(b)** was set at the median AUC in separate analysis using LPM as a reference line. The results are summarized from 50 replications.

For the identification of SNPs associated with two and three traits, the comparison performance of LPM and GPA are shown in Supplementary Fig. S11 and Supplementary Fig. S12, respectively. LPM performed better in terms of FDR control and AUC. In the identification of risk SNPs for both P1 and P4, a larger AUC can be achieved by integrating traits correlated with either P1 and P4, i.e., integrating P2, P3, P5 or P6.

When functional annotations do not play a role, we have shown that the relationship test graphs are similar for LPM, GPA and graph-GPA. In this case, we also compared their performance in the identification of risk SNPs of one specific trait. The results are shown in Supplementary Figs S13-S20. The performance of LPM and GPA is very close in terms of FDR and AUC. However, for graph-GPA, the empirical FDRs are conservative and AUCs are relatively small.

### 3.3 Simulations: type I error rate and power for the hypothesis testing of annotation enrichment

We further conducted simulations to evaluate the type I error rate and power of LPM for the hypothesis testing of annotation enrichment. We conducted simulations which considered only two traits with number of annotations *D* = 1 and correlation coefficient *ρ* = 0, and set the signal strength of the traits to be the same, i.e., *α*_1_ = *α*_2_ = *α*. We varied *α* in {0.2, 0.4, 0.6} to obtain the type I error rate, and varied the coefficient of annotation *β* in {−0.4, −0.3, −0.2, −0.1, 0.1, 0.2, 0.3, 0.4} to obtain the power of LPM for the enrichment test of the annotation. The results are shown in Supplementary Fig S21. We observed that the type I error rate was indeed controlled at the nominal level and the power was close to one when the signal strength was relatively strong (i.e., *α* = 0.2 or 0.4), and the coefficient was not very small (i.e., |*β*| ≥ 0.2).

### 3.4 Simulations: computational time

Figure 5 shows the computational time of LPM with *M* = 100, 000 and *D* = 5. For one pair of traits, the computational time depends on the signal strength of GWAS data and their correlation. When the number of traits increases, the time can be largely shortened by using more cores in parallel computation. However, the time is not linear in the number of cores. One reason is that the time we recorded not only included the time to fit bLPM for pairs of GWASs which is implemented parallelly, but also included the time for data preparation. Another reason is that we used Armadillo (30) in our R package which has already executed many functions (e.g., matrix multiplication) in parallel.

**Figure 5:**
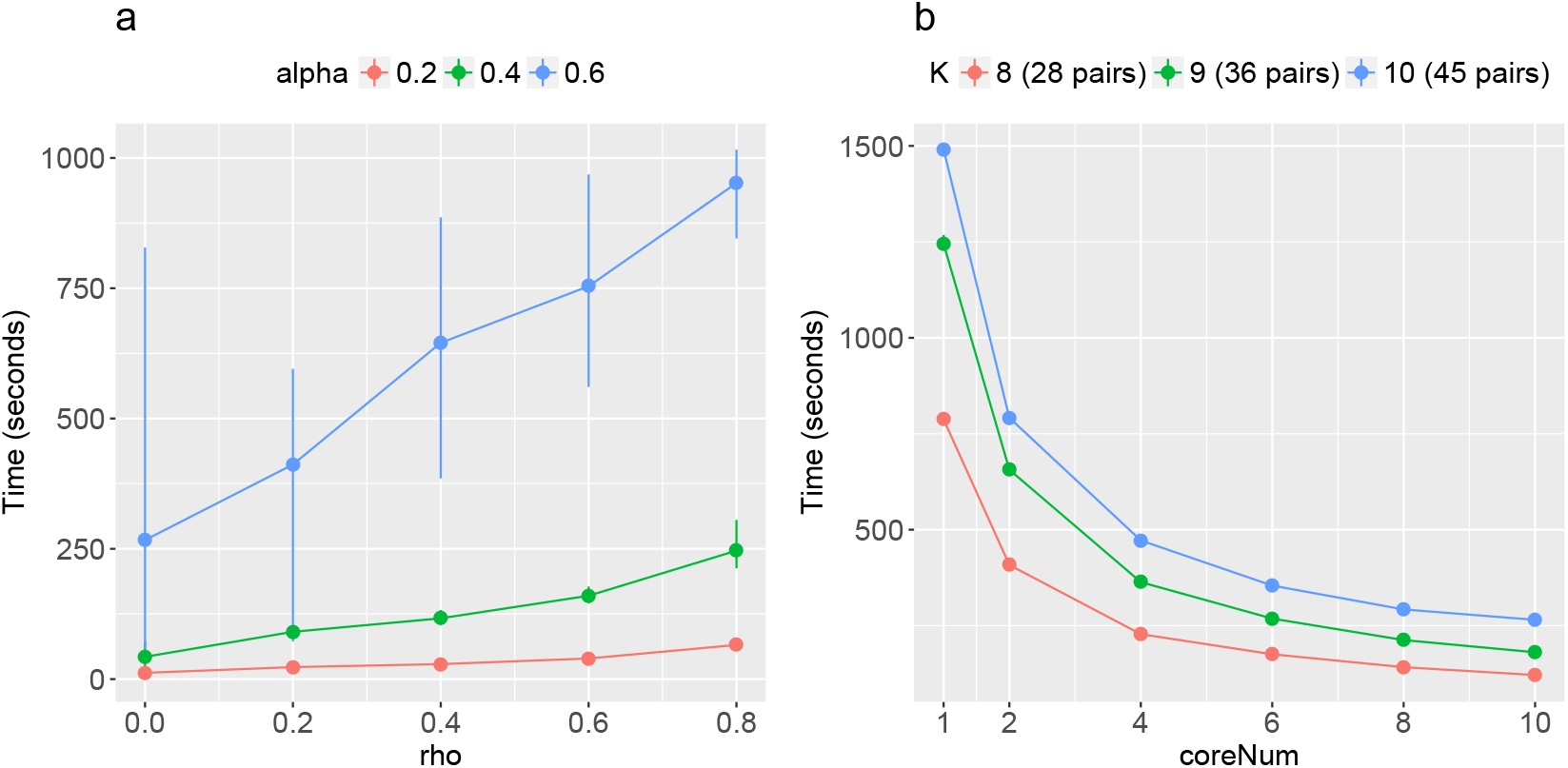
Computational time of LPM. **(a)** For one pair of traits, we varied the signal strength *α* and the correlation *ρ*. **(b)** For different numbers of traits, we varied the number of cores we used. The results are summarized from 10 replications.

### 3.5 Real data applications: correlations among 44 GWASs

We applied LPM to analyze 44 GWASs of complex traits integrated with 9 genic category annotations and 127 cell-type specific functional annotations (Materials and methods). The estimated correlations among 44 GWASs are shown in Figure 6. We observed that the correlations among traits were quite dense indicating that pleiotropy was pervasive (see Supplementary Table S3).

**Figure 6:**
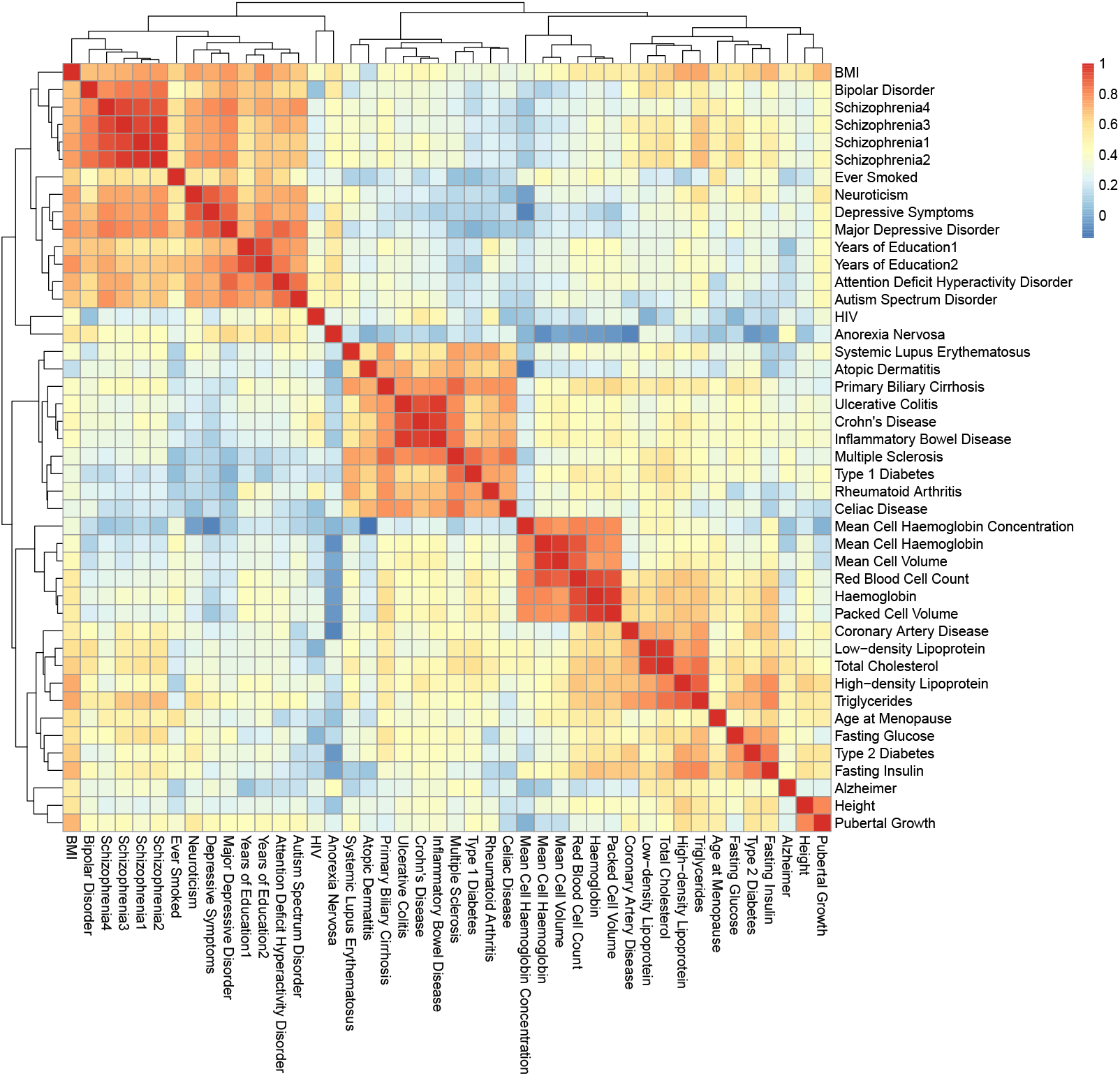
The estimated **R̂** for 44 GWAS with 9 genic category annotations and 127 cell-type specific functional annotations integrated.

According to Figure 6, traits can be divided into several groups with relatively strong correlations and these are consistent with the categories of traits. Main groups are psychiatric disorders (average of the estimated correlations in this group *ρ̄* = 0.7269) which include bipolar disorder (BIP) (31), schizophrenia (SCZ) (32, 33, 34, 35), neuroticism (36), depressive symptoms (36), major depressive disorder (MDD) (37), attention deficit hyperactivity disorder (38), autism spectrum disorder (39) and anorexia nervosa (40); hematopoietic traits (*ρ̄* = 0.8498) which include mean cell haemoglobin concentration, mean cell haemoglobin, mean cell volume, red blood cell count, haemoglobin and packed cell volume (41); autoimmune diseases (*ρ̄* = 0.7317) which include systemic lupus erythematosus (42), atopic dermatitis (43), primary biliary cirrhosis (PBC) (44), Crohn’s disease (CD) (45), ulcerative colitis (UC) (45), inflammatory bowel disease (45), celiac disease (46), rheumatoid arthritis (RA) (47), multiple sclerosis (48) and type 1 diabetes (T1D) (49); and lipid-related traits (*ρ̄* = 0.8427) which include high-density lipoprotein (HDL), triglycerides, low-density lipoprotein (LDL), and total cholesterol (TC) (50).

We also found some significant relationships between complex diseases and metabolic traits. For example, relatively high correlations were observed between coronary artery disease (CAD) (51) and lipid-related traits (*ρ̄* = 0.7929), among type 2 diabetes (T2D) (52), fasting glucose (53) and fasting insulin (53) (*ρ̄* = 0.7745). We observed that psychiatric disorders were correlated with many other traits, such as BMI (54) (*ρ̄* = 0.7242), years of education (55, 56) (*ρ̄* = 0.7167), HIV (57) (*ρ̄* = 0.6549) and ever smoked (58) (*ρ̄* = 0.6921). Similar evidence of the relationship has been found by Hartwig *et al.* (59), Breslau *et al.* (60), Chandra *et al.* (61) and Laurence *et al.* (62). We also discovered connections between height (63) and pubertal growth (64) *ρ̂* = 0.8111), between age at menopause (65) and fasting insulin (*ρ̂* = 0.6214), between Alzheimer (66) and lipid-related traits (*ρ̄* = 0.7109).

As a comparison, we used cross-trait LD Score regression to estimate the genic correlations among several traits (six hematopoietic traits, four lipid-related traits, CAD, height and pubertal growth). The results are shown in Supplementary Fig. S38. Although the definitions of correlation in cross-trait LD Score regression and LPM are different, the trends of relationship among these traits are quite similar because the two versions of correlation essentially capture the dependence among traits. We also applied graph-GPA to infer the relationship among traits. As graph-GPA was not scalable to a large number of traits, we can only analyze a subset of the traits (SCZ, CD, UC, RA, T2D, HDL, BIP and MDD). The relationship estimated by graph-GPA changed a lot as the number of traits changed (see Supplementary Fig. S39). This was because graph-GPA represented the relationship from a conditional independent structure, and the conditional independent structure might change when adding more traits or removing some included traits.

### 3.6 Real data applications: association mapping

We compared the number of SNPs identified to be associated with each of the 44 traits using LPM by five different analysis approaches: (i) separate analysis without annotation, (ii) separate analysis with genic category annotations, (iii) separate analysis with all annotations (genic category annotations and cell-type specific annotations), (iv) joint analysis of the top 1 correlated trait with all annotations and (v) joint analysis of the top 2 correlated traits with all annotations. The details of the top correlated traits are given in Supplementary Table S4. Using the fifth approach as a reference, we calculated the ratio of the number of risk SNPs identified using each approach.

Figure 7 shows that more risk SNPs can be identified by integrating functional annotations and correlated traits. For HIV, age at menopause and Alzheimer, a clear improvement was observed between the first two approaches, reflecting a significant enrichment of genic category annotations. Comparing the second and the third approaches, the contributions of cell-type specific functional annotations were observed to be significant for many traits, such as atopic dermatitis, fasting insulin, multiple sclerosis, PBC, RA and T1D. The differences between the fourth and fifth approaches were due to pleiotropy. For example, LDL and TC were observed to be highly correlated *ρ̂* = 0.9720). As a result, joint analysis of these two traits led to an improvement in identifying risk SNPs. Specifically, for the identification of risk SNPs of LDL with all annotations integrated, 3,758 SNPs were identified in separate analysis of LDL, whereas in joint analysis of LDL and TC, 7,845 SNPs were identified, when the global FDR was controlled at 0.1. The Manhattan plots are provided in Supplementary Fig. S40.

**Figure 7:**
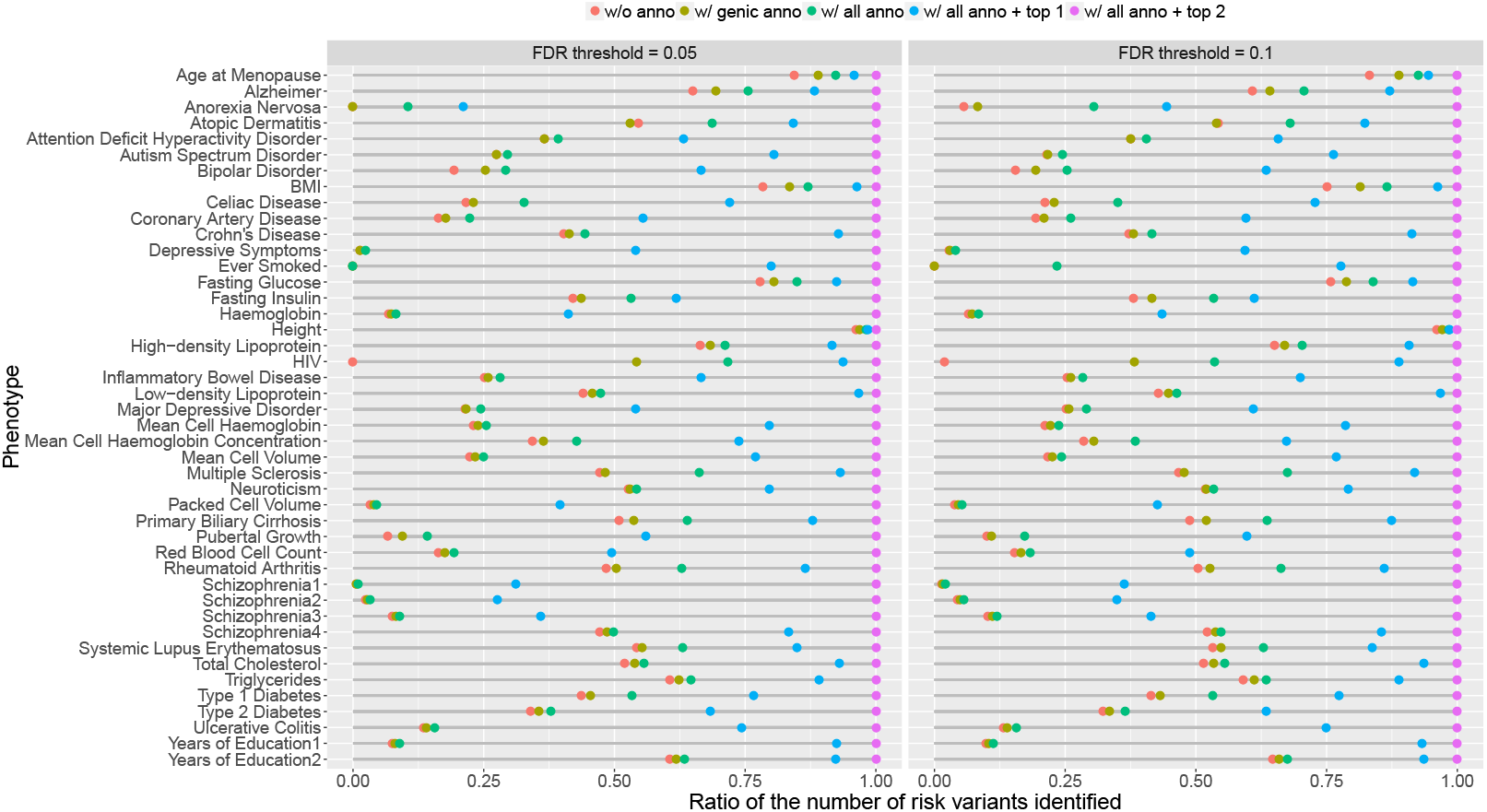
The numbers of variants identified to be associated with each of the 44 traits using LPM by five different analysis approaches: (i) separate analysis without annotation, (ii) separate analysis with genic category annotations, (iii) separate analysis with all annotations, (iv) joint analysis of the top 1 correlated trait with all annotations and (v) joint analysis of the top 2 correlated traits with all annotations. We controlled global FDR at 0.1. For visualization purpose, these numbers are normalized by dividing the corresponding number of variants identified by the fifth approach (joint analysis of the top 2 correlated traits with all annotations).

### 3.7 Real data applications: enrichment of functional annotations

The results for the enrichment test of nine genic category annotations and 127 cell-type specific functional annotations are shown in Figure 8. The detailed results of enrichment test are given in Supplementary Figs S41-S47 and the estimated coefficients of genic category annotations and cell-type specific functional annotation are given in Supplementary Figs S48-S52 and Figs S53-S59, respectively.

**Figure 8:**
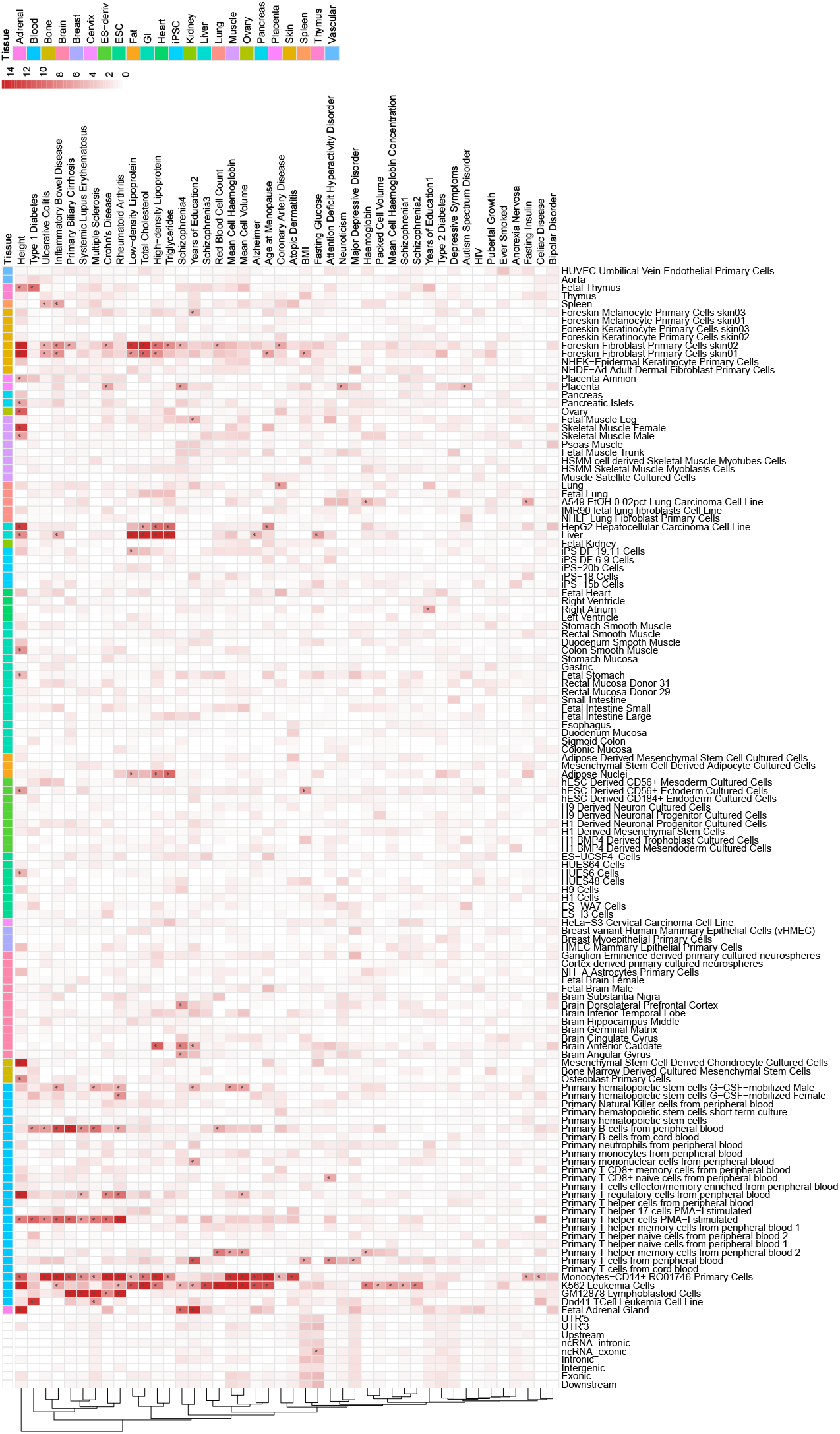
The −log_10_(*P*-value) for the enrichment test of nine genic category annotations and 127 cell-type specific functional annotations. The *P*-values which are smaller than 10^−15^ are set to be 10^−15^. The symbol “*” means the *P*-value is significant after Bonferroni correction at level 0.05.

We noted that the estimated coefficients for genic category annotations were positive for most traits except for intergenic. The estimated coefficients of the intergenic annotation were negative for many traits, e.g., *β̂* = −1.6649 (se = 0.9995) for puberal growth and *β̂* = −0.5992 (se = 0.7783) for ever smoked. Comparing the number of traits with significant coefficients for each genic category annotation, we found that exonic and UTR’3 were enriched most. Our results are consistent with the findings in the previous study (12).

For cell-type specific functional annotations, we detected enrichment of functional annotations in liver (liver and HepG2 cells) and fat (adipose nuclei) for lipid-related traits. Specifically, the enrichment in liver was significant for all the four lipid-related traits (HDL: *β̂* = 0.3015, se = 0.0256; LDL: *β̂* = 0.3609, se = 0.0267; TC: *β̂* = 0.3823, se = 0.0237; Triglycerides: *β̂* = 0.2478, se = 0.0293), which was consistent with findings in previous studies (16, 25, 17). SNPs annotated in cells of immune system were observed to be enriched for many traits, including autoimmune diseases, lipid-related traits, hematopoietic traits, some psychiatric disorders such as SCZ and BIP. For height, significant functional annotations included cells in immune system, bone, liver, muscle and skin. The foreskin fibroblast primary cells were shown to be enriched for some autoimmune diseases and lipid-related traits (17).

### 3.8 Real data applications: replications

Among the 44 GWASs, we analyzed four different GWASs of SCZ, Schizophrenia1 (9,379 cases and 7,736 controls) (32), Schizophrenia2 (9,394 cases and 12,462 controls) (33), Schizophrenia3 (13,833 cases and 18,310 controls) (34) and Schizophrenia4 (36,989 cases and 113,075 controls) (35). We found that the correlations among them were very high (*ρ̄* = 0.9505) and their enrichment of genic category annotations (see Supplementary Figs S48-S52) and cell-type specific functional annotations (see Supplementary Figs S60-S61) were highly consistent, indicating that the findings of LPM is replicable. As the sample size for GWASs from Schizophrenia1 to Schizophrenia4 became larger, the standard error of the estimated coefficients became smaller and the *P*-values of the enrichment test for annotations became more and more significant. For Schizophrenia1-3, only SNPs annotated in K562 Leukemia cells were enriched (Schizophrenia1: *β̂* = 0.1775, se = 0.0366; Schizophrenia2: *β̂* = 0.1846, se = 0.0310; Schizophrenia3: *β̂* = 0.1882, se = 0.0257). For Schizophrenia4, besides K562 Leukemia cells (*β̂* = 0.0830, se = 0.0178) more enrichment in functional annotations was detected, such as brain anterior caudate (*β̂* = 0.1076, se = 0.0179) and brain angular gyrus (*β̂* = 0.1208, se = 0.0238). As shown in Figure 7, more risk SNPs were identified to be associated with schizophrenia by jointly analyzing these GWASs. Similar results can be found (see Supplementary Figs S62-S63) for GWASs of educational traits, where Years of Education2 has a larger sample size than Years of Education1.

We also compared the results of UC, CD with IBD, and depressive symptoms with MDD. As IBD is comprised of two major disorders: UC and CD, depressive symptoms includes MDD, the consistent results from analyzing these GWASs also indicate the replicability of our method (see Supplementary Figs S64-S67).

## 4 Discussion

From the perspective of statistical modeling, a remarkable benefit of our model is its scalability to a large number of GWASs. Under LPM, the number of parameters in **R** only increases quadratically (*K*^2^) with the number of traits *K*. However, for other methods such as GPA, the number of groups will increase exponentially (2^*K*^). In particular, the design of LPM naturally allows a parallel algorithm such that the model fitting can be computationally efficient. The feasibility of the approach is demonstrated by our simulations in which the accuracy of parameter estimation using LPM is evaluated. Supplementary Fig. S22 shows that LPM provides satisfactory estimate of ***α***, ***β*** and **R**. Specifically, the estimates of ***α̃***, ***β̃*** and **R̃** using bLPM for different pairs are stable which shows the consistency and reliability of our algorithm (Figs S23-S32 in Supplementary). It also has theoretical supports which are based on the composite likelihood approach. The details of related theorem is provided in Supplementary Section S8. Because of the pairwise analysis, the number of SNPs for each pair of GWASs can be different. There is no need to remove SNPs with missing values in any one of the GWASs, avoiding the huge information loss especially for large amounts of traits.

LPM assumes that the *P*-values of SNPs in each GWAS are from a mixture of uniform and Beta distributions. We have shown that LPM is robust to violation of this assumption. We considered the situations when *P*-values in non-null group are from distributions other than the Beta distribution and when *P*-values are obtained from individual-level data (details are given in Supplementary Section S6.11 and S6.12 respectively). The results show that the type I error rate of LPM for the relationship test and the empirical FDR for identification of risk SNPs are well controlled at the nominal level.

In LPM, SNPs are assumed to be conditionally independent given the functional annotations. This assumption greatly simplifies our model and facilitates the computation and inference. However, in real application the genotype of SNPs are correlated in the presence of linkage disequilibrium (LD) effects. We conducted simulations to evaluate the impact of LD effects on our LPM model. The details of the simulations are given in Supplementary Section S6.13. The results (Supplementary Fig. S37) indicate that LPM can provide a satisfactory FDR control in terms of identifying a local genomic region of the risk SNPs.

In summary, we have presented a statistical approach, LPM, to integrate summary statistics from multiple GWASs and functional annotations. This unified framework can characterize relationship among complex traits, increase the statistical power for association mapping, integrate and investigate the effect of functional annotations simultaneously. With extensive simulations and real data analysis of 44 GWASs, we have demonstrated the statistical efficiency and computational scalability of LPM.

